# Literature Evidence in Open Targets – a target validation platform

**DOI:** 10.1101/124719

**Authors:** Şenay Kafkas, Ian Dunham, Johanna McEntyre

## Abstract

**Background:** We present the Europe PMC literature component of Open Targets – a target validation platform that integrates various evidence to aid drug target identification and validation. The component identifies target-disease associations in documents and ranks the documents based on their confidence from the Europe PMC literature database, by using rules utilising expert-provided heuristic information and serves the platform regularly with the up-to-date data since December, 2015.

**Results:** Currently, there are a total number of 1168365 distinct target-disease associations text mined from >26 million PubMed abstracts and >1.2 million Open Access full text articles. Our comparative analyses on the current available evidence data in the platform revealed that 850179 of these associations are exclusively identified by literature mining.

**Conclusion:** This component helps the platform’s users by providing the most relevant literature hits for a given target and disease. The text mining evidence along with the other types of evidence can be explored visually through https://www.targetvalidation.org and all the evidence data is available for download in json format from https://www.targetvalidation.org/downloads/data.

## 1 BACKGROUND

Understanding the underlying mechanisms of diseases is crucial in translational research. Discovering the association between drug target and disease has become a main focus for scientists since it is key for developing new drugs or repurposing them. Scientists gather various evidence representing different aspects of target-disease associations such as gene expression changes and the role of genetic variations to increase understanding. Such evidence can be stored in structured databases and requires integration to obtain complete and comprehensive knowledge in target validation studies.

Motivated by this, the Target Validation Platform (https://targetvalidation.org) [1] integrates different evidence from various resources with the aim of assisting scientists to identify and prioritise drug targets (proteins and their genes) associated with diseases and phenotypes. The evidence includes common disease genetic evidence based on GWAS study results from GWAS Catalog [2], rare Mendelian disease evidence based on ClinVar [3] clinical variant information from EVA and text mined target-disease associations from the Europe PMC (https://europepmc.org/) literature database [4] (see Table 3 for a complete list of evidence types).

Europe PMC contains over 33 million records and expands at a rate of over a million articles per year – one article every two minutes as scientists publish their findings continuously. Text mining target-disease associations is crucial for an integrated platform like the Target Validation Platform, since it provides a high volume of complementary and up-to-date data to the other type of evidences, otherwise the knowledge would stay hidden in millions of documents.

In this study, we present the Europe PMC Open Targets literature component that identifies target-disease associations in documents and ranks the documents according to their confidence based on rules utilising expert-provided heuristic information. Our main aim is to provide a scalable, robust and continuous text-mining service to the community for a real-world and very important application – target validation. Many of the previous studies focused on extracting gene-disease association from the literature [5-7]. However, only a few of them specifically focused on developing methods for integrated resources; DisGeNET [8] and DISEASES [9] for example cover various types of evidence for target validation. These two systems score target-disease associations for a given disease or target for their confidence extracted from MEDLINE abstracts and don’t provide very regular updates to the data. In DisGeNET, the target-disease text mining method is based on a machine learning approach while in DISEASES, target-disease associations are extracted based on scoring their co-occurrences according to their confidence. In comparison to DisGeNET and DISEASES, our system operates on full text articles in addition to abstracts, and ranks documents according to the confidence for a given target-disease association rather than the association itself. The confidence score of a given target-disease association is handled at the platform level and calculated based on all the evidence data in the platform by using a harmonic sum approach (see [1] for the details). Our approach to target-disease extraction differs from these systems, and probably many other traditional text-mining studies, in that we rely on heuristic information from experts/users for developing the system. The platform was first launched in December, 2015 and is publicly available at https://targetvalidation.org. Since then, our system has served the platform regularly (monthly) with up-to-date data.

## 2 METHODS

### 2.1 Resources Used

The literature source that we used in the study is the Europe PMC database. Europe PMC is one of the largest biomedical literature databases in the world which provides public access to >30.4 million abstracts and >3.3 million full text articles from PubMed and PubMed Central. In our analyses, we used the latest version of the Open Access full text articles (http://europepmc.org/ftp/archive/v.2016.06/) (~1.2 Million), and all of the PubMed abstracts (~26 Million) from the database.

Two comprehensive resources, UniProt and the Experimental Factor Ontology (EFO) are used to identify target and disease names in text, respectively. Two dictionaries are generated and refined from the human part of the SwissProt Database (the annotated part of UniProt, Release 2015_10) (http://www.uniprot.org/) and disease and phenotype parts of EFO (http://www.ebi.ac.uk/efo/) (Release 2.74) before applying text mining. In the refining process, we filtered out the terms that would introduce potentially very high numbers of false positives. These are the terms having character length < 3 (e.g. “A” is a gene name) and terms that are ambiguous with common English words (e.g. “Large” is a protein name as well). In addition, we generated term variations by replacing the widely used Greek letters in gene/disease names with their symbols (e.g. replacing “alpha” with α). The final target and disease dictionaries consisted of a total of 104,434 and 29,846 terms respectively. These dictionaries are available from ftp://ftp.ebi.ac.uk/pub/databases/pmc/otar/.

### 2.2 Target and Disease Name Annotation

We used the Europe PMC text-mining pipeline, which is based on Whatizit [10], to annotate target and disease names in text with the two dictionaries described above. Although we reduce a very high level of ambiguity by applying the dictionary refinement process before text mining the documents, some target and disease name abbreviations could still be ambiguous with some other names. For example, ALS which is an abbreviation used for “Amyotrophic Lateral Sclerosis”, is ambiguous with “Advanced Life Support” in some articles (e.g. see PMID:26811420). Therefore, we implemented and used a disease and target name abbreviation filter for screening out the potential false positive abbreviations introduced during the annotation process.

The abbreviation filter operates based on several rules using heuristic information. Regular expressions are used for identifying the text sequences in the form of “**X…˙. Y…. Z…. (XYZ)**”. The text in parentheses (i.e. (XYZ)) is identified as a gene/disease name abbreviation candidate if it is in the uppercase form, has length <6 (the length was decided by manually analysing a random subset of the Uniprot and EFO dictionaries) and annotated by the system either as a disease or a gene name, whereas, the text located immediately before the parentheses is identified as the potential long form. For example, in the following sentence from the article having PMID:26811420; “The guidelines form the basis for all levels of resuscitation training, now from first aid to *advanced life support (ALS)*,” the italicised text matches with our pattern defined above. “ALS” would be the abbreviation candidate and “advanced life support” would be the potential long form. Documents matching the pattern above are analysed manually by an expert to come up with heuristics that we can apply in filtering the ambiguous abbreviation. Abbreviation candidates satisfying one of the following rules are kept as true target/disease abbreviations, otherwise, they are filtered out:

For disease name abbreviation candidates:

- If any of the EFO long forms of the abbreviation candidate exists in the document
- If the long form extracted from the text contains any of the keywords (disease, disorder, syndrome, defect, etc.) that can be used to describe a disease

For gene or protein name abbreviation candidates:

- If (XYZ) appears more than 3 times in the document body (this rule applies to OA full text documents only)
- If the long form matches any of the terms from SwissProt or Enzymes (http://enzyme.expasy.org/)
- If the long form ends with (-ase/-ases) OR it contains any of the keywords (factor, receptor, gene, protein etc.) that can be used to describe a target name
- If at least 3 sentences for full text and at least 2 sentences for abstracts contain one of the keywords: “mutation, SNP, variation, gene, inhibit, variation, variant, polymorphism, mutant, isoform, protein, enzyme, activate, antibody, transcription, tumor suppressor, express, overexpress, regulator, receptor, oncogene” along with the protein name abbreviation candidate and a disease name.

### 2.3 Target-Disease Association Identification

Our association extraction method is based on identification of target-disease co-occurrences at the sentence level and applying several filtering rules to reduce noise possibly introduced by the high sensitivity, low specificity co-occurrence method. Our filtering rules utilise heuristic information from a careful manual analysis of the text data to filter out potential false positive associations. More specifically, the manual analyses are conducted iteratively by analysing a randomly selected set of results and identifying the reasons behind the false positives in the results so that we could formulate them as filtering rules to tune our system.

The system applies the following filtering rules:

1. Filter out all type of articles except “Research” articles (e.g. Reviews, Case Reports).
2. Filter out target-disease associations appearing in the following sections: Methods, References, Acknowledgement & Funding, Competing Interests, Author Contribution and Supplementary Material.
3. Filter out target-disease associations that appear only once in the body of a given article but not in the article’s title or abstract.

Sections of a given document are identified by using our Section Tagger [11] tool that we developed previously.

### 2.4 Document Scoring

A document scoring algorithm is implemented and integrated in to the system to assign each document a confidence score for a given target-disease association. Document confidence scores are used to rank all the documents relevant to a given target-disease association. The algorithm is based on weighting document sections and sentence locations differently for full text articles and abstracts respectively (see Table 1 and Table 2). The weighting approach is often used in text mining tasks for assigning confidence scores. For example in [9] different weights are assigned to the different features for calculating the confidence scores of the identified associations. In our study, we assign weights from the range of [1-10] which is wide enough to pick different weights for different sections based on their potential confidence. The following formulas, CS_1_ and CS_2_ are used to calculate the confidence scores for abstracts and full text articles respectively:

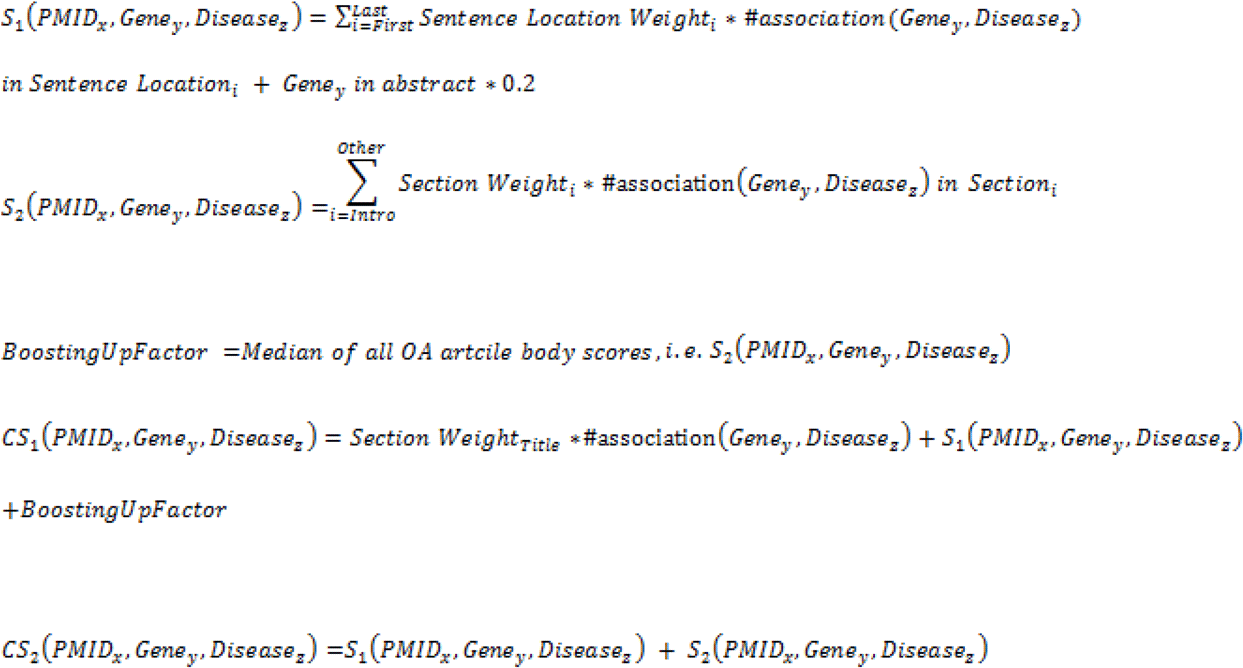

**Table1:** Sentence location weights in abstracts.

| Sentence Location | Weight |
| --- | --- |
| First or second | 2 |
| Last | 5 |
| Other | 3 |

**Table 2:** Section weights in full text articles.

| Section | Weight |
| --- | --- |
| Title | 10 |
| Abstract | See Table 1 |
| Results, Figure, Table | 5 |
| Discussion, Conclusion | 2 |
| Introduction, Case Study, Appendix, Other | 1 |

The weights are selected based on heuristic information and our goal is to identify associations that are the subject of the given paper, rather than instances that are reviewing prior knowledge. Therefore, we assign the highest weight, which is 10, to “Title”, since an article title would contain the most confident information and highlight the main finding of the study. The lowest weight (1), is assigned to “Introduction”, since well-known associations are often reported here while a higher weight (5) is assigned to the “Results”, “Figures” and “Tables” sections where the new findings are generally reported.

The sentence location weights that are used for abstract scoring are determined based on a sentence level concept analysis by using CoreSC [12]. CoreSC is a text-mining tool which assigns each sentence one of its 11 pre-defined concepts such as “Results” and “Background”. Our concept analysis performed on randomly selected 360 MEDLINE abstracts revealed that most of the time, the last sentence of a given abstract is a “Results” sentence, while the first/second one is generally an introductory sentence (“Background”) (CoreSC analysis results are available at ftp://ftp.ebi.ac.uk/pub/databases/pmc/otar/). We further verified our finding by manually checking some of the abstracts from this set. Hence, we assign the highest weight (5) to the last sentence and lower weights to the first/second and other sentences accordingly.

## 3 RESULTS & DISCUSSION

### 3.1 Performance Evaluation

The ultimate goal of this study is to provide a scalable, robust and continuous service to the biomedical community for target validation, by using text mining methods. Therefore, we took a different approach from many traditional text mining studies and benchmarked the system based on expert perspective - expert satisfaction and feedback are the most valuable parameters for us to judge on the system’s performance. Our service has been up and running since December 2005 and we continuously improve our algorithms as we receive user feedback. Nevertheless, as a case study, we estimated the overall performance of the system on two randomly selected samples by using Mean Average Precision (MAP) which is a commonly used metric in evaluation of ranking system performance. MAP takes into account the relative order of the documents retrieved by the system and gives more weight to the documents returned at higher ranks [13]. We manually estimated the MAP for abstracts only as 89% and for full text articles as 90% on the top 25 documents of the two randomly selected gene-disease associations which were IGF1 – Diabetes and NOD2 – Inflammatory Bowel Disease. We also estimated the correlation coefficients between the abstract only and full text article scores as 0.82 and 0.94 for IGF1 – Diabetes and NOD2 – Inflammatory Bowel Disease respectively. Obtaining almost the same MAP values for both abstracts only and full text articles as well as high correlation coefficients between the scores confirm our heuristic score adjustment.

The abbreviation name filtering performance alone was estimated to have an F-Score value of 92.3% by evaluating randomly selected 50 sentences from the Open Access articles reporting on target-disease associations.

### 3.2 Added value from the literature mined target-disease associations

The Target Validation Platform currently covers evidence from literature mining, genetic associations, somatic mutations, known drugs, gene expression, affected pathways and animal models. (Please refer to [1] for further information about how the other types of evidence data are gathered.) In the current release (release 1.2) of the platform, there are a total number of 2,485,000 distinct target-disease associations. Table 3 shows a comparison of the target-disease association data currently available in the platform. The literature evidence constitutes the largest amount of data compared to the other type of evidence (such as gene expression and animal models). Currently, there are more than 1.1 million (47% of the whole evidence data) distinct target-disease associations extracted from ~26 million PubMed abstracts and ~1.2 million open access full text articles. Other large amounts of evidence data are provided from the gene expression (~900K) and animal models (~600K) sources. The analysis shows that 21.75% (197943) of gene expression, 43.31% (56228) of genetic associations, 69.36% (2506) of affected pathways, 16.55% (99836) of animal models, 33.59% (19801) of somatic mutations and 34.56% (19811) of known drugs evidence data overlap with the literature mining data. The majority of the distinct associations in the platform are identified exclusively through literature mining (~850K, 34.21%) showing the added value from text mining.

**Table 3:** Comparison on the target-disease association data in the Target Validation Platform (release 1.2)

| Evidence Type | Total number of distinct target-disease associations | Overlapping target-disease association |  |  |  |  |  | Total number of exclusively identified associations |
| --- | --- | --- | --- | --- | --- | --- | --- | --- |
|  |  | Gene Expression | Genetic Associations | Affected Pathways | Animal Models | Somatic Mutations | Known Drugs |  |
| Literature Mining | 1168365 | 197943 | 56228 | 2506 | 99836 | 19801 | 19811 | 850179 |
| Gene Expression | 909960 | X | 18945 | 901 | 35616 | 32795 | 9913 | 669330 |
| Genetic Associations | 129826 | X | X | 1912 | 26504 | 3626 | 2133 | 62999 |
| Affected Pathways | 3613 | X | X | X | 1045 | 310 | 163 | 714 |
| Animal Models | 602995 | X | X | X | X | 2965 | 4421 | 486167 |
| Somatic Mutations | 58941 | X | X | X | X | X | 1845 | 16197 |
| Known Drugs | 57319 | X | X | X | X | X | X | 33005 |

Total number of distinct target-disease associations in the platform is 2485000.

The discrepancy between the literature mining data and the other type of evidence data is due to the fact that each evidence data is gathered by using different methods as well as resources. For example, gene expression data is gathered from Expression Atlas (https://www.ebi.ac.uk/gxa/home), the scope of which is microarray or RNA-Seq experiments. Other evidence data such as genetic associations and known drugs are gathered through manual curation of the literature by experts and from DailyMed (https://dailymed.nlm.nih.gov/dailymed/). Our approach is based on computationally extracting evidence data from the literature. In many of the curated studies, which may report associations between many targets and several diseases, it is unusual to highlight the individual association results in a way that is detectable by the sentence co-occurrence approach and often these associations are confined to a supplementary data table. Although text mining and manual curation both use the biomedical literature as a resource, the coverage of the methods is different and complementary. In fact, in our early work with users the text-mining approach was highly valued precisely because it accesses evidence from papers that do not contribute to the curated databases. One further reason for any discrepancy originates from the licencing restrictions on the reuse of full text content. We can only text mine the full text of Open Access publications (and all MEDLINE abstracts), while experts can curate evidence from the non-open access publications, accessed for reading via journal subscriptions.

We further analysed the contribution of text mining based on the associations by disease and associations by target in Table 4 and Table 5 respectively. Table 4 shows comparison of the associations by disease in the platform. Currently, there are a total number of 9426 associations by disease in the platform. The majority of these diseases are provided from genetic associations (5912), literature mining (5801) and animal models (4942). Our analysis shows that 56.02% (405) of gene expression, 59.98% (3546) of genetic associations, 88.89% (504) of affected pathways, 68.86% (3403) of animal models, 53.75% (494) of somatic mutations and 82.72% (1489) of known drugs provided target associated diseases overlap with the literature mining data. The majority of the distinct associations by disease in the platform are identified exclusively through genetic associations (1336, 14.17%) and literature mining (1304, 13.83%).

**Table 4:** Comparison of the associations by disease in the Target Validation Platform (release 1.2)

| Evidence Type | Total number of distinct associations by disease | Overlapping associations by disease |  |  |  |  |  | Total number of exclusively identified associations by disease |
| --- | --- | --- | --- | --- | --- | --- | --- | --- |
|  |  | Gene Expression | Genetic Associations | Affected Pathways | Animal Models | Somatic Mutations | Known Drugs |  |
| Literature Mining | 5801 | 405 | 3546 | 504 | 3403 | 494 | 1489 | 1304 |
| Gene Expression | 723 | X | 520 | 196 | 309 | 460 | 328 | 25 |
| Genetic Associations | 5912 | X | X | 527 | 3725 | 530 | 1193 | 1336 |
| Pathways | 567 | X | X | X | 443 | 168 | 310 | 9 |
| Animal Models | 4<br>942 | X | X | X | X | 281 | 752 | 811 |
| Somatic Mutations | 919 | X | X | X | X | X | 354 | 113 |
| Known Drugs | 1800 | X | X | X | X | X | X | 179 |

Total number of distinct associations by disease in the platform is 9426

**Table 5:** Comparison of the associations by target data in the Target Validation Platform (release 1.2)

| Evidence Type | Total number of associations by target | Overlapping associations by target |  |  |  |  |  | Total number of exclusively identified associations by target |
| --- | --- | --- | --- | --- | --- | --- | --- | --- |
|  |  | Gene Expression | Genetic Associations | Affected Pathways | Animal Models | Somatic Mutations | Known Drugs |  |
| Literature Mining | 14728 | 14217 | 8670 | 664 | 5187 | 3903 | 736 | 321 |
| Gene Expression | 29842 | X | 9817 | 671 | 5449 | 4125 | 743 | 14148 |
| Genetic Associations | 10200 | X | X | 561 | 4072 | 3165 | 569 | 217 |
| Pathways | 690 | X | X | X | 379 | 324 | 70 | 4 |
| Animal Models | 5497 | X | X | X | X | 3744 | 484 | 8 |
| Somatic Mutations | 4138 | X | X | X | X | X | 330 | 2 |
| Known Drugs | 756 | X | X | X | X | X | X | 1 |

Total number of distinct associations by target in the platform is 30592

Table 5 shows comparison of the associations by target in the platform. Currently, there are a total number of 30592 associations by target in the platform. The majority of these targets are provided from gene expression (29842), literature mining (14728) and genetic associations (10200). Our analysis shows that 47.64% (14217) of gene expression, 85% (8670) of genetic associations, 96.23% (664) of affected pathways, 94.36% (5187) of animal models, 94.32% (3903) of somatic mutations and 97.35% (736) of known drugs provided disease associated targets overlap with the literature mining data. The majority of the distinct associations by target in the platform are identified exclusively through gene expression (14148, 46.25%) which is understandable given the comprehensive gene coverage in gene expression experiments such as RNA-seq.

Altogether, our analysis shows that literature mining suggests many more new target-disease associations (850,179, see Table 3) rather than new diseases (1304, see Table 4) or targets (321, see Table 5) involved in associations.

### 3.3 Examples of target-disease associations exclusively identified by literature mining

Our analysis reveals that there are a total number of 850,179 target-disease associations exclusively identified by literature mining. One such example is the CTGF gene and male breast carcinoma association (Figure 1) (https://www.targetvalidation.org/evidence/ENSG00000118523/EFO_0006861). Currently, there is evidence for the association of 101 different targets with male breast carcinoma. All of these targets are identified through literature mining and only 4 of them are also supported by the known drugs evidence.

**Figure 1:**
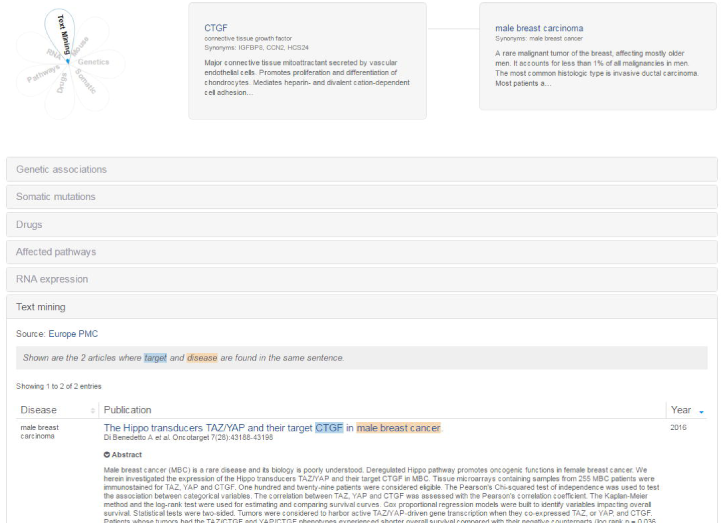
The CTGF and male breast carcinoma association.

Another example is the ST3GAL4 and diabetes mellitus association. There are 1572 different publications potentially reporting this association (Figure 2). (https://www.targetvalidation.org/evidence/ENSG00000110080/EFO_0000400). Currently, there is evidence for the association of 5017 different targets with diabetes mellitus. 3670 of these targets are identified through literature mining.

**Figure 2:**
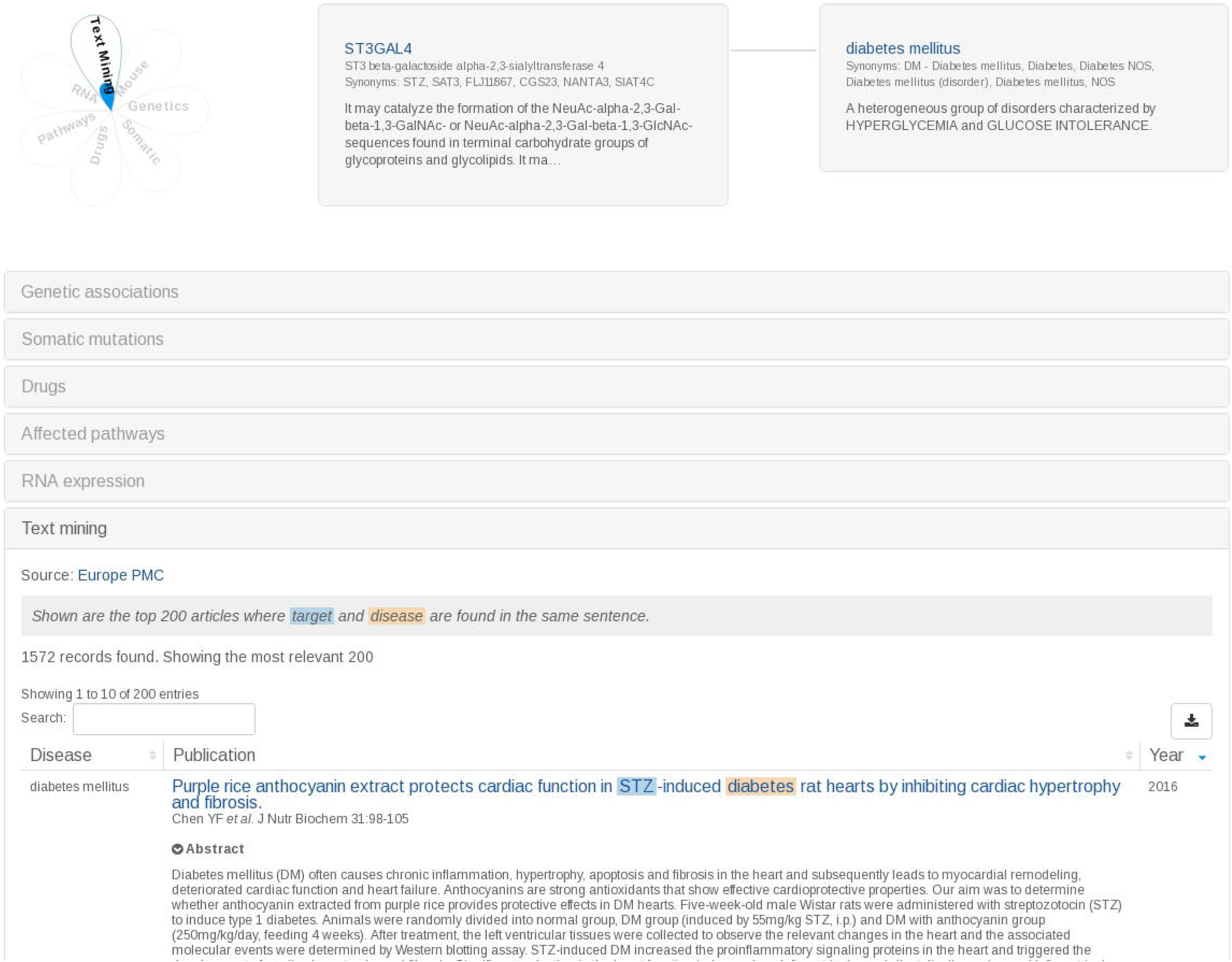
The ST3GAL4 and diabetes mellitus association.

### 3.4 User Experience

Since the first release of the Europe PMC Open Targets component, we iteratively improved our text mining algorithm and the visualisation of the text mining evidence in the Target Validation Platform based on user feedback. Initial user testing showed that the incorporation of the text mining evidence in to the platform filled in perceived gaps in evidence caused by limitations in coverage by the other direct evidence sources. The users also valued the reinforcement of other evidence when complementary text mining evidence was available. Feedback from users of incorrect associations predominantly from false positive entity recognition assisted us in improving our filters.

## 4 CONCLUSIONS

Here, we present the Europe PMC Open Targets component, a new service for analysing and visualising target-disease associations from the literature within Open Targets. The aim of this component is to help users by providing the most relevant literature hits for a given target and disease. The platform users reported that the text mining evidence helped Open Targets to become more complete and a given association is more credible when it is supported not only by text mining but also by the other types of evidence. Our text mining algorithm and visualisation of the text mining evidence are improved iteratively based on user feedback.

Currently, we are analysing the EFO coverage by comparing it against the other existing disease/phenotype resources such as Disease Ontology (http://disease-ontology.org/) and Unified Medical Language System (https://www.nlm.nih.gov/research/umls/). In future, we plan to expand the EFO’s coverage based on our findings. We also work on classifying articles based on the available evidence types in the platform such as genetic variations and RNA expression. This would provide users with a better understanding and more insight on the weight of individual target-disease associations.

## List of Abbreviations

EFO: Experimental Factor Ontology
Europe PMC: Europe PubMed Central
PMID: PubMed Identifier
CS: Confidence Score
MAP: Mean Average Precision
RNA: Ribonucleic Acids
GSK: GlaxoSmithKline

## Declarations

### Availability of data and material

All target-disease data is available for download from https://www.targetvalidation.org/downloads/data as compressed json files.

The compiled target and disease dictionaries as well the dataset used in MAP estimation are available from ftp://ftp.ebi.ac.uk/pub/databases/pmc/otar/ for download.

### Authors’ contributions

ŞK, ID and JM conceived of the study. ŞK implemented the software and performed the experiments. ID provided the heuristic information and performed the manual evaluations. All authors evaluated the results and contributed to the manuscript. All authors read and approved the final manuscript.

## Acknowledgements

We thank Maria Liakata for her help with running CoreSC on a sample set of abstracts. This paper is an extended version of a conference paper by the same name, presented at Phenotype Day@ ISMB 2016.

